# Mechanical Regulation of the Cytotoxic Activity of Natural Killer Cells

**DOI:** 10.1101/2020.03.02.972984

**Authors:** L. Mordechay, G. Le Saux, A. Edri, U. Hadad, A. Porgador, M. Schvartzman

## Abstract

Mechanosensing has been recently explored for T cells and B cells and is believed to be part of their activation mechanism. Here, we explore the mechanosensing of the third type of lymphocytes – Natural Killer (NK) cells, by showing that they modulate their immune activity in response to changes in the stiffness of a stimulating surface. Interestingly, we found that this immune response is bell-shaped, and peaks for a stiffness of a few hundreds of kPa. This bell-shape behavior was observed only for surfaces functionalized with the activating ligand MHC class I polypeptide-related sequence A (MICA), but not for control surfaces lacking immunoactive functionalities. We found that stiffness does not affect uniformly all cells but increases the size of a little group of extra-active cells, which in turn contribute to the overall activation effect of the entire cell population. We further imaged the clustering of costimulatory adapter protein DAP10 on NK cell membrane and found that it shows the same bell –shape dependence to surface stiffness. Based on these findings, we propose a catch-bond-based model for the mechanoregulation of NK cell cytotoxic activity, through interaction of NKG2D activating receptors with MICA. Our findings reveal what seems to be “the tip of iceberg” of mechanosensation of NK cells, and provides an important insight on the mechanism of their immune signaling.

**Statement of Significance:** The mechanical sensing of immune lymphocytes was recently demonstrated for T cells and B cells, but not for the third type of lymphocytes – Natural Killer (NK) cells. Interestingly, previous reports on lymphocyte mechanosensing were controversial, and showed either positive or negative changes in their immune activity with environmental stiffness, depending on the stiffness range. In this paper, we directly demonstrated that NK cells modulate their response with the stiffness of the stimulating surface, and this modulation has a bell-shape trend. We found that there is a strong correlation between the response to stiffness and clustering of adaptor proteins. Upon this correlation, we proposed a mechanosensing model based on the catch-bond nature of activating ligand-receptor complexes in NK cells.

## Introduction

Cells use mechanical forces to probe their environment. In the last two decades, mechanosensing and mechanotransduction have been extensively studied, mostly in the context of focal adhesions, and they are known today to play key roles in many vital cell functions, such as adhesion, proliferation, mitosis, motility and death (1–5). In addition, emerging studies have shed light onto the role of mechanical forces in the function of immune cells (6). Mechanistic features of the immunological synapse – a functional interface between a lymphocyte and an antigen presenting cell (APC) (7, 8) – stems from actin retrograde flow which drives the adhesion molecules and antigen receptors toward the central zone of the immunological synapse (9, 10). Furthermore, it has become progressively clear that the antigen receptors themselves, such as T cell receptors (TCRs) and B cell receptors (BCRs), have mechanosensing and mechanotransducive characteristics (11, 12).

The growing evidence for the mechanical aspects of lymphocyte signaling greatly supports the emerging paradigm that these cells use mechanical forces to discriminate between healthy and damaged cells (13). This paradigm has been clearly demonstrated in a few recent *in vitro* studies, showing that lymphocytes functionally respond to variations in the stiffness of the surface they contact. However, the conclusions of these studies were controversial. Some studies consistently showed an increase in the immune response, including immune activation of T cells and B cells with increasing surface stiffness (14–17), indicating that these two cell types might share similar mechanosensing features. The substrate stiffness in these studies, however, was mostly limited to a scale of tens to hundreds kPa. Conversely, a decrease in T cell activation with surface stiffness was reported for a stiffness range of 100kPa to ∼ 2MPa (18). Could it then be possible that the immune response of lymphocytes to surface stiffness is bell-shaped? This question cannot be firmly addressed based on the compilation of all above-mentioned studies, since they used cell-stimulating surfaces of different materials, different activation ligands, cells of different sources, and different experimental conditions. Each of these factors produces its own effect on cell stimulation. Hence, to address this question, the immune response of cells to the environmental stiffness must be studied for the broadest possible range of stiffnesses, whereas all the other parameters are kept identical. Yet, as of today, such a study has not been performed, to the best of our knowledge.

Whereas the mechanical aspects of immune signaling have been studied mostly in T cells and B cells, much less is known about the role of mechanical forces in the function of a third type of lymphocytes – Natural Killer (NK) cells. NK cells are the sentinels of the innate immune system, and they contribute to the immune protection by direct cytotoxicity, cytokine secretion, and the regulation of adaptive responses of T-cells. The immune response of NK cells is regulated through a delicate balance of multiple activating and inhibitory signals (19–21). Early indication that mechanical forces also play a role in the regulation of NK cell immune activity was recently provided by *Barda-Saad and Co*, who showed that actin retrograde flow controls NK cell immune response *via* conformational changes of tyrosine phosphatase SHP-1 (22). Shortly after, we reported that NK cells stimulated onto vertical nanowires functionalized with activating ligands produced an enhanced immune response (23). In that study, the nanowires – nanomechanical objects with ultra-high aspect ratio and flexibility – exhibited extraordinary compliance to cellular forces as small as 10pN. We thus proposed that this high compliance produced a strong mechanical stimulus, which resulted in a much stronger degranulation compared to that of cells stimulated on a flat rigid surface. Still, the comparison between nanowires and rigid surfaces represents two extreme cases. The nature of mechanical interactions between NK cells and their environment should still be systematically studied for the widest possible range of mechanical and biochemical stimuli, individually and in combination with each other.

In this work, we carried out the first, to the best of our knowledge, systematic study of the immune response of NK cells to the stiffness of their environment. For this purpose, we stimulated primary NK cells onto elastic surfaces whose stiffness varied from 50kPa to 3MPa. This range, which spans over two orders of magnitude, encompasses most of the ranges previously used to study T cells and B cells. In order to provide NK cells with biochemical stimulation, we functionalized these surfaces with MHC class I polypeptide-related sequence A (MICA) molecules that are specifically recognized by the NK cell activating receptor NKG2D (24). We assessed the immune response of NK cells to the surface stiffness *via* surface expression of lysosomal-associated membrane protein-1 (CD107a), and found that it was non-linear, and had a bell shape characteristic with a peak at 200kPa. To separately assess the effect of each stimulus – mechanical and chemical – we also stimulated NK cells on control surfaces that had no bioactive functionalities and found that these control surfaces yielded relatively low degrees of activation, whose dependence on the surface stiffness was negligible. We also quantified the spreading of the stimulated NK cells, and found that it does not correlate with their immune activity. Concurrently, we found that the surface stiffness of 200kPa stimulated the formation of the highest number of clusters of DAP10 – an adaptor protein of the NKG2D receptor, indicating that the enhanced activation for this surface stiffness is associated with NKG2D organization and signaling. Overall, these results clearly demonstrate the existence and importance of mechanosensing in NK cells, and provide an intriguing insight into their non-linear response to the physical features of their environment.

## Materials and Methods

### Sample Fabrication and Biofunctionalization

Elastic surfaces of polydimethylsiloxane (PDMS) were prepared using a standard elastomer kit Sylgard 184 (Dow Inc.), with different hardener-resin ratios of 1:50, 1:35 and 1:10 (w/w) (25). The mixture was poured onto a glass slide and smeared, to achieve PDMS thickness of ∼0.3mm (26). Finally, the PDMS samples were degassed in a desiccator, and cured for one hour at different temperatures: 130°C for 1:20 and 1:35 ratios, and 200°C for 1:10 ratio (27). Elastic modulus was assessed form force-distance curves of contact mode AFM (Asylum Research, MFP-3D-Bio, Hertz model (17, 28, 29), see SI for details). To functionalize PDMS surfaces with MICA ligands, the samples were first treated in Oxygen plasma for 5 seconds (Harrick PDC-32G), and then modified with (3-Aminopropyl) triethoxysilane (APTES) by immersing into 10% APTES solution in ethanol for 10 minutes, rinsing with ethanol and drying for 10 minutes in oven at 80°C (30). Then, the samples were incubated overnight at 4°C in 2μg/mL with solution of His-MICA (SinoBiological) in Phosphorous Buffer Solution (PBS).

### Immunofluorescence characterization of PDMS functionalization

Fluorescence microscopy was used to confirm that MICA was efficiently immobilized onto APTES-functionalized PDMS. For this purpose, MICA was labelled with a red fluorophore as follows. A 1:50 v /v of 5(6) carboxytetramethyl-rhodamine N-succinimidyl ester (TAMRA-NHS) was added to 10 μl MICA with 5μl NaHCO_3_ and 34μl of PBS. The mixture was left to react for one hour. The mixture was purified using a protein purification kit (Life technologies). APTES-functionalized PDMS was then immersed in the TAMRA-MICA solution overnight at 4°C in 2μg/mL. Afterwards, we then rinsed the samples twice with PBS, and once in pure water, and mounted them on a coverslip with DAKO Fluorescence Mounting Medium (Agilent). The samples were imaged with an epifluorescence microscope (Nikon Eclipse Ti2-E). Please see supporting information for more details.

### Isolation of Primary NK Cell (pNK)

Cells were isolated using a human negative selection-based NK isolation kit (RosetteSep, STEMCELL Technologies). Purified NK cells were then cultured in stem cell serum-free growth medium (CellGenix GMP SCGM, 20802-0500) supplemented with 10% heat-inactivated human AB plasma from healthy donors (SIGMA, male AB, H-4522), 1% l-glutamine, 1% Pen-Strep, 1% sodium pyruvate, 1% MEM-Eagle, 1% HEPES 1M, and 300 IU/mL recombinant human IL-2 (PeproTech). pNK cells were purified from peripheral blood of healthy, adult, volunteer donors, recruited by written informed consent, as approved by the Institutional Review Board Ben-Gurion University of the Negev (BGU).

### NK Cell Activation Experiments

Cultured pNK cells were seeded onto the surfaces in an amount of 0.5 million cells per sample in fresh medium (containing 50U/ml IL2) prepared from SCGM and RPMI (1:6 v/v) supplemented with APC anti-human CD107a (1:1000 v/v). The cells were incubated for 3 hours on the surfaces at 37°C. After the incubation, the surfaces were rinsed once in PBS, fixed with 4% paraformaldehyde (PFA), rinsed once again PBS, and then directly stained with Alexa Fluor 555 phalloidin (Life Technologies) diluted 1:40 v/v in blocking buffer (5% skim milk in PBS) without permeabilization for one hour in 37°C to prevent damage to the cell membrane. Finally, the samples were rinsed once with PBS and once in pure water, then nuclei were stained by mounting the samples on coverslips with Fluoroshield Mounting Medium with DAPI (Abcam) to prepare the samples for imaging. NK cells were imaged using a Nikon eclipse Ti2-E epifluorescence microscope system (20x Ph1 DLL air objective, wavelengths of 365, 550 and 635 nm for DAPI, Alexa Fluor 555 and CD107a respectively). Please see supporting information for details on the methodology used to quantify CD107a expressions.

### Imaging DAP10 clustering

pNK cells were seeded on the surface for 10 minutes and fixed according to the protocol described in the previous section, excluding the anti-CD107a supplement.

After permeabilization (Triton X-100, 0.1% in PBS, 15 min at 4 °C), the cells were incubated 1 hr. at 37 °C in Alexa Fluor 488 phalloidin (Life Technologies, 1:40 v/v in blocking buffer) and Alexa Fluor 594 labelled anti-human DAP10/HCST (R&D Systems, 1:500 v/v in blocking buffer) to stain for the cytoskeleton and DAP10 clusters respectively. Finally, the samples were rinsed once with PBS and once in pure water, then the nuclei were stained by mounting the samples on coverslips with Fluoroshield Mounting Medium with DAPI (Abcam) to prepare the samples for imaging. DAP10 clusters were characterized using a Zeiss LSM 880 confocal microscope system equipped with the Airyscan detector for super-resolution microscopy (31) (63x 1.4 NA DIC M27 oil objective, wavelengths of 405, 488 and 561 nm). Please see supporting information for more details.

## Results and Discussion

### NK cells produce bell-shape response to the substrate stiffness

In order to find whether and how NK cells modulate their immune activation in response to environmental stiffness, we prepared elastic surfaces of polydimethylsiloxane (PDMS) with variable elastic moduli – 50kPa, 200kPa, and 3MPa. We experimentally confirmed the elastic moduli of the obtained PDMS substrates using AFM (Fig. S1). To add a biochemical stimulus to the PDMS surfaces, we functionalized the PDMS with MICA ligands. For this purpose, we first functionalized the PDMS with APTES, which facilitates the physisorption of biomolecules *via* electrostatic interactions (32).

We confirmed that MICA was efficiently immobilized onto APTES-functionalized PDMS by incubating these surfaces in an aqueous solution of TAMRA-labelled MICA, and imaged the obtained surfaces using an epifluorescence microscope. The strong contrast in fluorescence emission between the TAMRA-MICA functionalized surface and APTES-functionalized surfaces (Fig. 1a,b) confirms the effective chemisorption of MICA. To ensure that the elastic modulus is the only parameter that is changed between different samples, and the stimulating conditions are not affected by the difference in the amount of MICA, we quantified the surface density of MICA molecules for each sample (see supporting information). To do so, we first assessed the fluorescent signal of a known quantity of TAMRA labeled MICA (Fig. S2a) and used it as a calibration factor to assess the surface density of MICA on our different surface stiffnesses of PDMS samples. We found that this surface density was quite similar for all the samples, ranging between 4000 to 8000 molecules per square micron (Fig. S2b). Given the fact that MICA has a diameter of about 12 nm (33), we can deduce that MICA was immobilized onto PDMS surface in almost closely packed configuration. Furthermore, the obtained density of MICA is far above the minimal density required for NK cell activation, which is about ∼100 molecules per square micron (34). Therefore, we created here a stimulation environment for NK cells with the maximum possible amount of biochemical stimulus, while the mechanical stimulus, which is defined by the surface stiffness, was the only variable.

**Figure 1:**
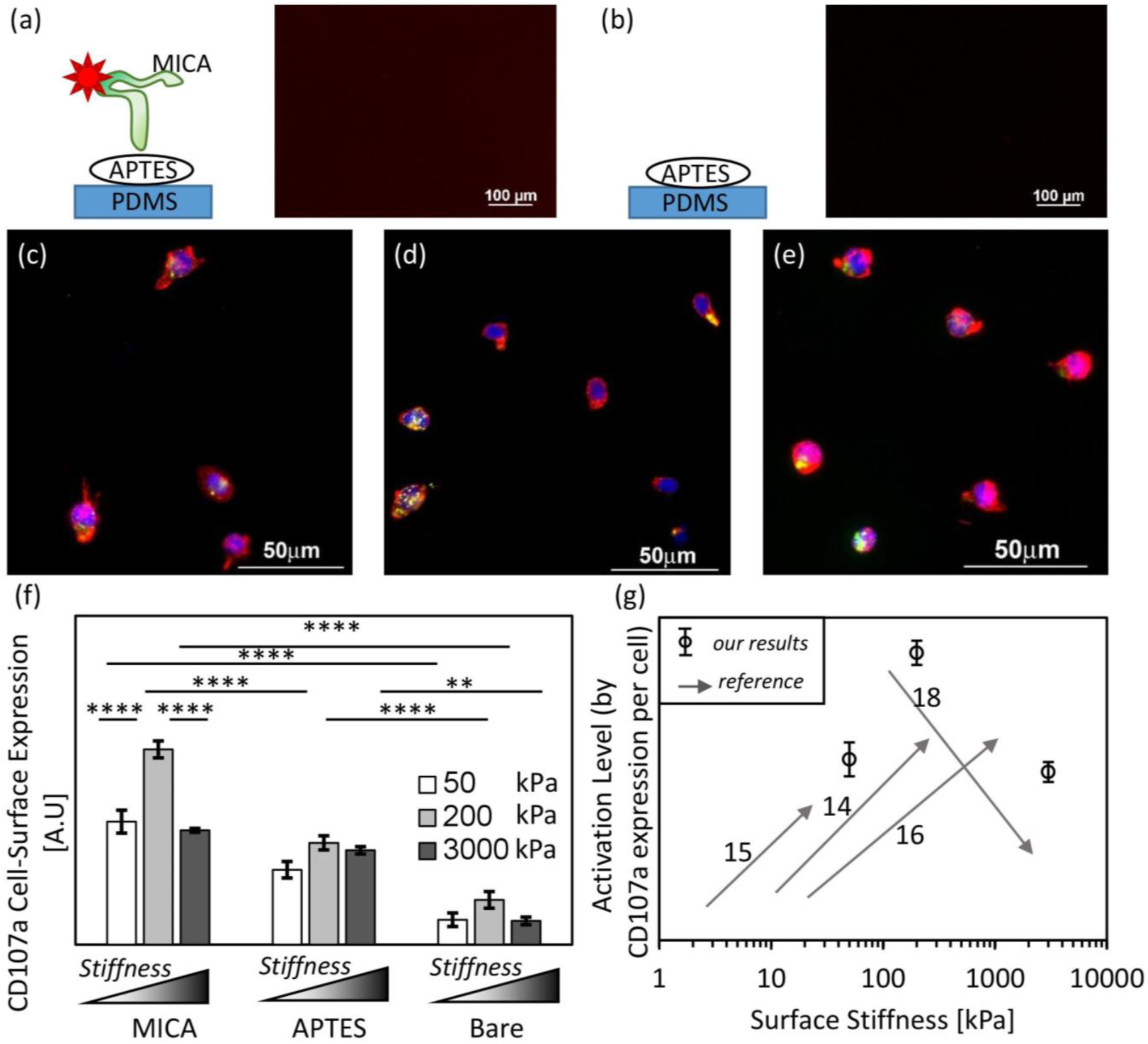
Fluorescence images of PDMS substrates functionalized with (a) APTES only and (b) APTES followed by TAMRA-labelled MICA. The pictures shown here have the contrast range. (c-e) Representative images of NK cells on MICA functionalized PDMS with stiffnesses of 50kPa, 200kPa and 3MPa, respectively. Cytoskeleton was stained with phalloidin (red), nucleus – with DAPI (blue), and CD107a with its fluorescently labeled mAb (green). (f) Degree of CD107a cell-surface expression of NK cells on MICA modified, APTES-modified and bare PDMS respectively. The degree of CD107a was quantified by summarize the fluorescence intensity of the APC-labeled anti-CD107a per cell (for ease of view, the APC labelled anti-CD107a was pseudo-colored in green in c-e). The results are shown the compilation of at least three different experiments and for at least three sets of samples per experiment. Using Graphpad Prism, analysis of variance and Tukey’s post hoc tests were performed to assess the significant changes in behavior – four stars (****) represent p<0.0001. (g) Compilation of activation trends reported for T cells and B cells is shown by arrows with the corresponding reference numbers, and our results for NK cell are shown in circles.

We incubated primary NK cells on the elastic surfaces for three hours. In order to elucidate separately the effect of chemical and mechanical stimulations on NK cell immune response, we used two types of control surfaces: PDMS functionalized with APTES but without MICA, and bare PDMS. We quantitatively assessed the degree of immune activation of NK cells by imaging the surface expression of the lysosome-associated membrane protein-1(CD107a), which is a common degranulation marker (35, 36). To that end, we added fluorescently labelled anti-CD107a to the incubation medium. After incubation, we fixed and stained the cells with phalloidin and DAPI for cytoskeleton and nucleus visualization, respectively. We quantified the activation degree of NK cells on each tested surface by measuring the average signal produced by the CD107a antibody which had accumulated at the cell surface.

Fig. 1c-e shows representative images of NK cells stimulated on MICA-functionalized PDMS with the moduli of 50kPa, 200KPa, and 3MPa, respectively. CD107a accumulated on the cell membrane and stained with its fluorescent antibody appears in bright green. Fig. 1f shows the average signal of CD107a per cell on MICA-functionalized surfaces and on the control surfaces. As expected, MICA-PDMS stimulated a significantly higher CD107a expression of NK cells compared to that obtained for control surfaces. Furthermore, CD107a expression on MICA-PDMS greatly varied with stiffness, and this variation was non-linear and bell-shaped, with a maximum CD107a signal for 200kPa. On the other hand, on the control surfaces, the effect of surface stiffness on the CD107a expression was significantly less pronounced. This result largely mirrors our previous report on the activation of NK cells on nanowires, where NK cells showed significant activation only on nanowires functionalized with MICA, while control surfaces lacking either nanowires, MICA, or both, did not induce a substantial activation of NK cells (23).

The observed bell-shape response is highly intriguing, and its exact mechanism is yet to be understood. Importantly, this is the first time, to the best of our knowledge, that the immune response of NK cells to variations in environmental stiffness is systematically studied. However, it could be of interest to compare the observed behavior of NK cells to that of other lymphocytes stimulated in analogous conditions. As mentioned above, the effect of surface stiffness on immune activation was previously reported for T cells and B cells, but those reports used relatively short ranges of surface stiffnesses, and showed different stiffness versus activation trends that appeared to depend on the studied stiffness range (14–18, 37) (Table 1). As mentioned, the combination of those studies for T cells suggested the possibility of a bell-shape correlation (6). While this possibility should be separately verified in the future for T cells and B cells, our result mirrors the state-of-the-art studies (Fig. 1g), and shows that at least for NK cells, the bell-shape immune response to stiffness indeed exists.

**Table 1:** State-of-The-Art. The existing studies which assess the influence of the surface stiffness on the immune activity.

| Reference | Cell Type | Stiffness Range<br>(kPa) | Activation with<br>Increase in Stiffness |
| --- | --- | --- | --- |
| *E. Judokusumo et al.* (2012) <sup>(14)</sup> | T | 10-200 | Increase |
| *Z. Wan et al.* (2013) <sup>(15)</sup> | B | 2.6-22.1 | Increase |
| *Z. Wan et al.* (2015) <sup>(16)</sup> | B | 20-1100 | Increase |
| *R.S. O'Connor et al.* (2012) <sup>(18)</sup> | T | 100-2300 | Decrease |

As anticipated, bare PDMS surfaces produced the lowest immune activation of NK cells, due to the absence of any chemical stimulus. Still, the effect of surface stiffness of bare PDMS is also bell shaped. These results highlight the need for a chemical stimulus to allow lymphocytes, such as NK cells, to mechanically sense their environment, and transduce this sensing into a functional immune response. In addition, APTES functionalized surfaces stimulated a higher immune response compared to bare PDMS. Here, the effect of surface stiffness was negligible. We presume that APTES stimulated NK cell activation due to the positive charges of the terminal amino groups, which produce asymmetric charge distribution across the cell membrane, and might positively affect NKG2D signaling similarly to that of TCR in T cells (38).

### Stiffness of antigen-functionalized surfaces regulates the adhesion of NK cells

We next focused on whether and how surface stiffness and biochemical functionalization affect the adhesion of NK cells. Importantly, our PDMS surfaces had no specific adhesion functionalities that can stimulate adhesion mediated by NK cell adhesion receptors, such as Intercellular Adhesion Molecule 1 (ICAM), but only MICA that engages NKG2D receptors. We estimated the degree of adhesion by counting the number of cells per unit area after mounting the samples for imaging (Fig. 2a). Adhesion, and its dependence on surface stiffness, varied between the probed surface chemistries. As expected, bare PDMS surfaces yielded the lowest degree of adhesion, due to a lack of any biochemical functionalities that specifically recognize NK cell receptors. Surprisingly, for MICA-functionalized surfaces we obtained a consistent increase in the degree of cell adhesion with an increase in surface stiffness. Conversely, the control surfaces (both bare and APTES) produced NK cell adhesion bell-shape tendency with increasing surface stiffness. The relatively high adhesion observed on APTES-PDMS can be explained, again, by its positive charges (39). Our results indicate that specific and non-specific functionalities signal *via* different pathways, and that their signals, while being integrated with the mechanical signals produced by the surface stiffness, affect differently cell adhesion. Although the mechanism of this integration has to be further studied, we can conclude at this stage that we established an experimental toolbox which can independently control the adhesion of NK cells *via* either biochemical stimulus produced by the chemical functionalities, or by mechanical stimulus produced by the surface modulus of elasticity.

**Figure 2:**
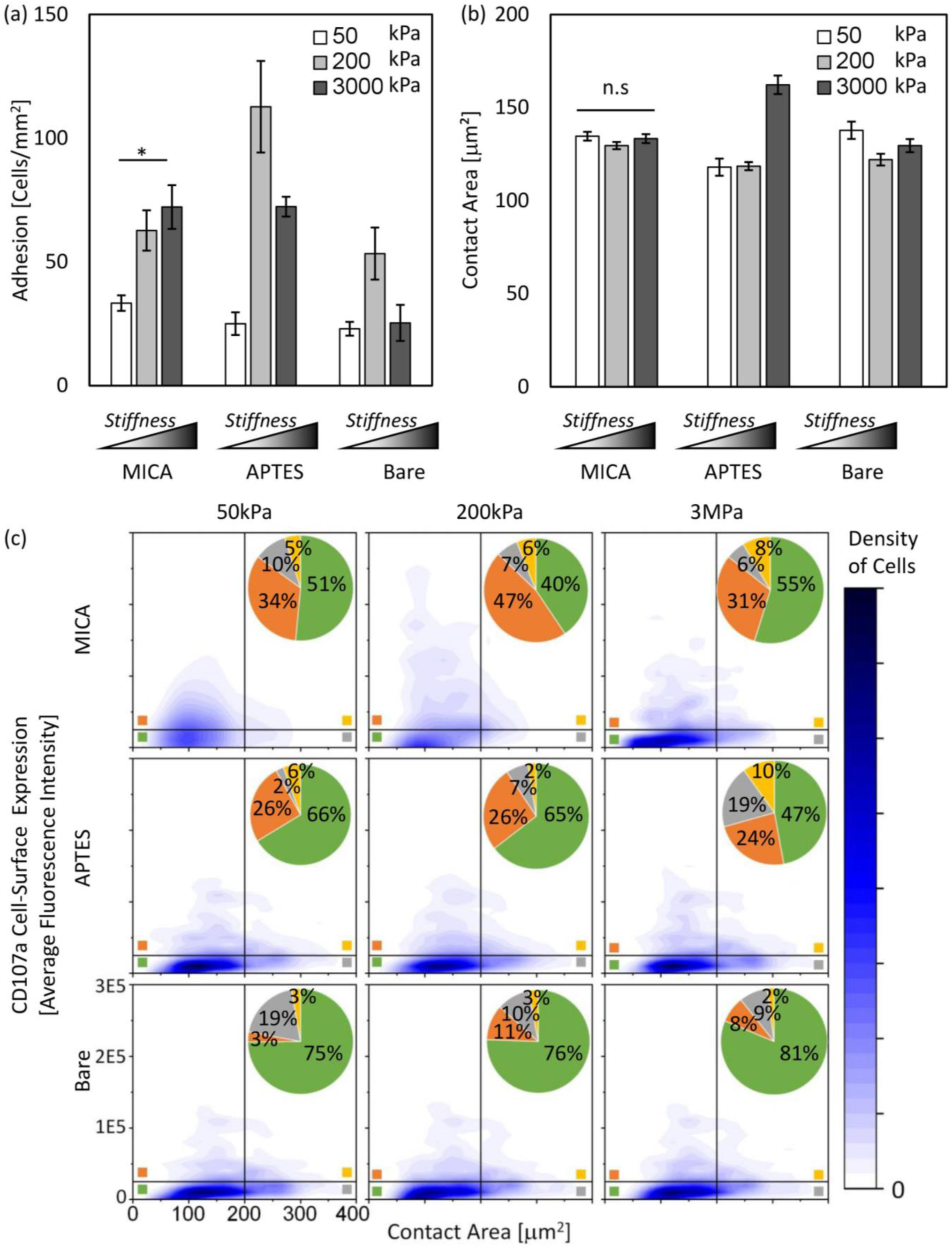
(a) NK cell adhesion vs. stiffness. Cell adhesion was calculated by counting the projected contact area of the cell cytoskeleton divided by the area of the surface that they adhere to, for each type of sample. (b) NK cell spreading vs. stiffness. Cell spreading was quantified by measuring the projected contact area of the cell cytoskeleton, for each type of sample. The results are shown the compilation of at least three different experiments and for at least three sets of samples per experiment. Using Graphpad Prism, analysis of variance and Tukey’s post hoc tests were performed to assess the significant changes in behavior – four stars (****) represent p<0.0001. (c) Density plots of spreading (defined by contact area) vs. activation (defined by degree of CD107a). Percentages of cells for each of the four sub-populations is shown in the inset. The results are shown the compilation of at least three different experiments and for at least three sets of samples per experiment. The plots are analyzed by using Origin software.

### Stiffness of antigen-functionalized surface does not affect NK cell spreading but regulates the number of highly activated NK cells

It was previously shown that the spreading of NK cells does not necessarily require a interaction with adhesion functionalities LFA-1, but can occur via MICA ligation, which, however, produces a cell morphology different than that induced by LFA-1 ligation (40). Furthermore, we recently showed that NK cell spreading depends on the spatial density of MICA ligands, and that at least 100 MICA ligands per square micron are required to stimulate full spreading of NK cells (34). Still, all the above studies were done on rigid glass surfaces coated with ligands, and the role of surface stiffness in spreading of NK cells has not been explored yet.

Here, we assessed the impact of stiffness on the spreading of NK cell, by measuring their projected area after 3 hours of incubation (Fig. 2b). We found that in the case of MICA functionalized PDMS, surface stiffness had no effect on NK cell spreading. This finding mirrors the previously reported lack of correlation between spreading and activation of NK cells (41). Conversely, APTES functionalization stimulated higher spreading on stiffer PDMS.

We also examined whether there is a correlation between spreading and activation degree. To this end, for each set of stimulation conditions (i.e. stiffness + chemistry) we divided cells into four sub-populations defined by spreading and activation gates. The values for the spreading and activation gates for the 2D spreading-activation density plots (Fig. 2c) were determined by the average + standard error of the cell areas and CD107a signals for bare PDMS. It can be seen from the 2D spreading-activation density plots (Fig. 2c), that for the majority of the stimulating conditions, the distribution between the populations is very similar, and thus the plots have very similar outlines. For these outlines, more than half of the cells fall into the category defined by low spreading and low activation, while a relatively low percentage of cells showed either high activation or high spreading, and a negligible percentage of cells showed both.

A clear exception in the spreading-activation plot is for 200kPa PDMS functionalized with MICA, which induced the highest average degree of degranulation, as shown in Fig. 1f. In this plot, we observe an exceptionally high fraction (>50%) of NK cells defined as “highly-activated” by our “activation threshold”. Surprisingly, the vast majority of these highly activated NK cells (47% out of a total of 54%) were less spread, i.e. below the “spreading threshold”. Indeed, *Culley et al*. recently demonstrated that the extent of spreading for NK cells is not proportional to the outcome of signaling balance in NK cells, but rather occurs if a threshold of activation is surpassed (40). This lack of direct proportionality between spreading and immune activation, especially when estimated after a relatively long time of incubation, is observed for all the combinations of surface elasticity and surface chemistry presented herein but is more pronounced for 200kPa PDMS functionalized with MICA.

What also follows from the spreading–activation plots of 200kPa PDMS functionalized with MICA, and from its comparison to other conditions, is that the high activation on 200kPa MICA-PDMS (Fig. 1f) stems not from a uniform increase in the activation in all the cells, but from an increase in the sub-population of highly activated cells, while the cells out of this sub-population retain their low activation level. Interestingly, this observation reflects recent work, in which individual NK cells placed in micro-wells together with target cells were categorized into sub-populations by their cytotoxic response, e.g. the number of killed target cells (41). It was found that there is a small but extremely active sub-population NK cells, which was called “serial killer” NK cells, due to their ability to eliminate a number of target cells each (42). This *in vitro* detection of the “serial-killer” population mirrors a prevalent model by which only a minority of NK cells is responsible for tumor elimination. The existence of sub-populations with different cytotoxic activities can be related to the fact that NKG2D expression varies amongst cells (Fig. S3). Thus, the highly active cells are presumably those expressing larger amounts of NKG2D. Whereas the percentage of extra-active NK cells was initially the same for all the tested samples, we showed that by tuning mechanical elasticity of the stimulating environment, the relative fraction of extra-active NK cells that undergo degranulation can be controllably manipulated, and the change in this fraction affects the average activation level of the entire population of NK cells. Therefore, mechanical signaling should not be viewed as a stimulating factor that can gradually increase the immune activity of each NK cell alone, but rather as a stimulating factor that helps more NK cells overcome a certain activation threshold.

### Effect of the surface stiffness on the activation of NK cells is associated with the receptor microclustering

Organization and clustering of receptors within the membrane of lymphocytes is the key factor in their activation (43, 44). The cytotoxic activity of NK cells is tightly regulated by the microclustering of their activating receptors such as CD16 (45) and NKG2D (46), as well as of their inhibitory receptors such as KIR2DL1 (47). This microclustering can be regulated in vitro by various micro-/nano-platforms with controlled spatial distribution of activating and inhibitory ligands(48–51). On the other hand, it was natural to ask whether surface stiffness controls the cytotoxic activity of NK cells by modulating the clustering of target receptors of the used activating ligands, or *via* a signaling pathway independent of receptor clustering. Since binding NKG2D to MICA could possibly prevent its availability for a fluorescent antibody, we did not stain directly NKG2D. Instead, we labeled its transmembrane adaptor DNAX-protein 10 (DAP10), whose hexametric complex with NKG2D activates signaling in NK cells (52, 53), and therefore has a proximity of few nanometers to the NKG2D receptor (54). Furthermore, colocalization of NKG2D and DAP10 clusters was recently demonstrated at the nanometric scale within the membrane of activated NK cells (54). Notably, receptor clustering is an early signaling event, and NK cells were reported to form mature DAP10 clusters at the cell membrane within a few minutes (54). Thus, to quantify DAP10 clustering, we imaged the cells after 10 minutes of incubation.

The clusters of DAP10 were imaged using a Zeiss LSM880 confocal microscope equipped with an Airyscan detector for super-resolution microscopy (31). We analyzed DAP10 clustering using ImageJ by identifying clusters and measured their density per cell and their size (see SI for details). Fig. 3a shows typical microclusters of DAP10 in NK cells stimulated on MICA-functionalized surface with stiffnesses of 50kPa, 200kPa, and 3MPa, and the quantification of cluster density and cluster size. We found that the total amount of DAP10 adaptors protein expressed by NK cells is similar for all the tested surfaces (Fig. 3b). On the other hand, we found that surface stiffness alters the way DAP10 receptors are clustered within the cell membrane. The highest density of clusters per cell and, concurrently smallest clusters were found in cells cultured on MICA functionalized PDMS with a stiffness of 200kPa (Fig. 3c,d). Notably, the density of clusters per cells *versus* surface stiffness mirrors the already familiar bell-shape behavior observed for NK cell activation, with a peak at 200kPa.

**Figure 3:**
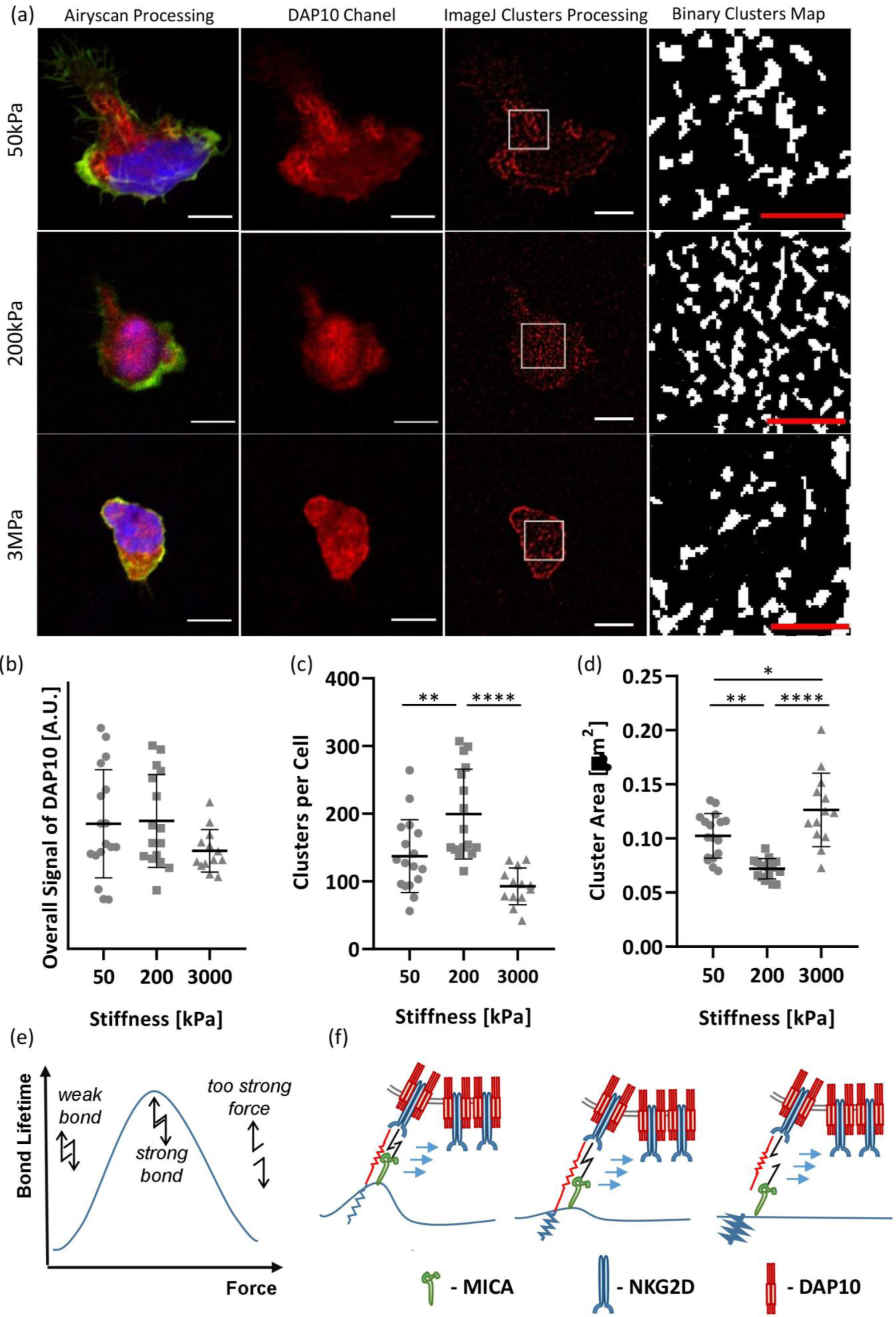
Clustering of NKG2D. (a) Representative images of NK cells on MICA functionalized PDMS with stiffnesses of 50kPa, 200kPa and 3MPa, respectively. Cytoskeleton was stained with phalloidin (green), nucleus – with DAPI (blue), and DAP10 with its fluorescently labeled (red). White scale bar-5μm, red scale bar-2μm. Clustering of NKG2D for each PDMS sample with stiffnesses of 50kPa, 200kPa and 3MPa are shown in (b) overall signal of DAP10 per cell (c) density of clusters per cell and (d) average cluster area per cell (d). The results are shown the compilation of at least three different experiments and for at least three sets of samples per experiment. Using Graphpad Prism, analysis of variance and Tukey’s post hoc tests were performed to assess the significant changes in behavior – four stars (****) represent p<0.0001. (e) Scheme of catch-bond lifetime vs. applied force. (f) Proposed model for the effect of surface stiffness on the receptor clustering.

These results strongly suggest a correlation between surface stiffness-regulated NK activation and the clustering of DAP10 adaptors, and most probably its associated activating receptors NKG2D, although the latter was not directly imaged. Indeed, the obtained correlation between the activation of NK cells and clustering of DAP10 mirrors similar correlation recently reported by *Balint et al.* (54). There, the effect of different ligands on the activation of NK cells was studied, and it was found that MICA stimulated the highest degranulation, but also the organization of NKG2D into more numerous and smaller clusters. Notably, in that work, the ligands were immobilized onto a rigid surface, and the biochemical stimulus provided by MICA was the only environmental parameter through which activation was controlled. Thus, while receptor clustering is undoubtedly the key factor in the regulation of NK cytotoxic activity, this factor itself can be regulated either by biochemically *via* antigens, or physically *via* mechanical properties of the environment. The final outcome, i.e. activation, is produced by the cumulative effects of the biochemical and physical stimuli.

The exact mechanism by which environmental stiffness regulates receptor clustering in NK cells, and therefore their activation, remains unclear. One scenario is based on the possibility that NKG2D receptors sense mechanical forces and transduce mechanical signals *via* conformational changes. Such a mechanosensing mechanism has been recently established for TCR receptors in T cells (55), and depicts the TCR-pMHC complex as a “catch-bond”, whose life-time increases with the applied force up to a peak of ∼10pN (56), and then decreases when the force is further increased. The catch-bond characteristics of the TCR-pMHC complex were explored experimentally using optical tweezers (57), micropipette based adhesion assays (58), and biomembrane force probe (56), as well as modeled in terms of TCR and membrane conformation (59). This catch-bond mechanism has also been observed in NK cells for the CD16 activating receptor (60), but not yet for NKG2D. Still, this scenario could explain the observed bell-shaped effect of surface stiffness on receptor clustering, and on the immune response of NK cells (Fig. 3e,f). Since MICA is electrostatically tethered to the elastic surface, both the surface and the MICA-NKG2D complex can be considered as two independent elastic components interconnected in series. While the MICA-NKG2D complex moves toward a cluster formed by other NKG2D molecules, each elastic component deforms, and the deformation of each component depends on their relative spring constants. If the surface is relatively soft (Fig. 3f left), it would contribute to most of the deformation, while the MICA-NKG2D complex would stay in its “loose” state. This scenario corresponds to the left part of the force versus life-time bell curve. With an increase in surface stiffness, the deformation between the two components are distributed more evenly, and the catch bond reaches its optimal conformation, and therefore, the longest lifetime, which corresponds to the peak of the curve (Fig. 3f middle). Further increase in surface stiffness induces a deformation that cannot be withstood by the bond, which eventually breaks (Fig. 3f right). Again, the observed functional response of NK cells to the variation of surface stiffness can be at most an indirect indication that MICA-NKG2D complex is indeed a catch-bond. The direct demonstration of the catch-bond behavior of MICA-NKG2D complex should be done to support the proposed model, similarly to the above-mentioned works on TCR-pMHC agonist catch-bond.

## Conclusion

In summary, our paper provides a new insight on the diversity of environmental stimuli for NK cells, and demonstrates their effects, separately and in combination with each other, on NK cell activation. Among these stimuli, the biochemical ones have been the most broadly studied so far. In contrast, the physical stimuli are still far from being fully explored. Here, we not only provided the first direct evidence that NK cells sense the stiffness of the surface they contact, but also revealed the trend with which this stiffness affects their immune response. This trend mirrors the compilation of recent studies on T cells and indicates that both types of cells have similar mechanosensing characteristics. The observed bell-shape response to surface stiffness shows that there is an optimal range of stiffness for NK cell stimulation, and NK cells likely utilize their ability to sense the stiffness in their decision making as to whether a target cell should be attacked or tolerated. The correlation between the bell-shaped immune response to stiffness and the bell-shaped dependence of receptor clustering indicates that these receptors are involved in mechanosensing, and probably act as mechanotransductors. The proposed catch-bond mechanism of MICA-NKG2D can explain the observed bell-shape response, but still needs to be verified. Finally, studies of physical signaling in lymphocytes focused so far mostly on the effect of the environmental stiffness. Yet, there has been increasing evidence of many additional physical factors that regulate the immune function of cells. For example, it has been recently shown that not only stiffness, but also the viscosity of the environment regulates cell function (61). Regarding cytotoxic lymphocytes such as T cells and NK cells, the combined effect of stiffness, viscosity, and additional physical factors on their immune function is still to be explored. Our work exposes what seems to be “the tip of iceberg” of the physical sensing of NK cells. This sense undoubtedly deserves further investigation, as it will help understand the mechanism of the immune activity of NK cells, which is both fundamentally important, and can facilitate the development of future immunotherapies.

## Author Contributions

L.M, G.L.S, A.P, and M.S designed the experiment, L.M and G.L.S performed the experiments, L.M, G.L.S, A.E, and U.H analyzed the data, all the authors contributed to the writing of the manuscript.

## Acknowledgments

This work was funded by the Multidisciplinary Research Grant - The Faculty of Health Science in Ben-Gurion University of the Negev and Israel Ministry of Science and Technology, Israel Science Foundation, Individual Grant #1401/15, and Israel Science Foundations: F.I.R.S.T. Individual Grant #2058/18. The manuscript was written through contributions from all authors.

